# Co-evolutionary analysis accurately predicts details of interactions between the Integrator complex subunits

**DOI:** 10.1101/696583

**Authors:** Bernard Fongang, Yingjie Zhu, Eric J. Wagner, Andrzej Kudlicki, Maga Rowicka

**Affiliations:** Department of Biochemistry and Molecular Biology, University of Texas Medical Branch, Galveston, TX 77555, USA; Glenn Biggs Institute for Alzheimer’s & Neurodegenerative Diseases, UT Health San Antonio, San Antonio, TX 78229, USA; Department of Biochemistry and Structural Biology, UT Health San Antonio, San Antonio, TX 78229, USA; Department of Epidemiology and Biostatistics, UT Health San Antonio, San Antonio, TX 78229, USA; Institute for Translational Sciences, University of Texas Medical Branch, Galveston, TX 77555, USA; Sealy Center for Molecular Medicine, University of Texas Medical Branch, Galveston, TX 77555, USA; Informatics Service Center, University of Texas Medical Branch, Galveston, TX 77555, USA

**Keywords:** residue coevolution, INTS4, INTS9, INTS11, Gaussian convolution

## Abstract

Solving the structure of large, multi-subunit complexes is difficult despite recent advances in cryoEM, due to remaining challenges to express and purify complex subunits. Computational approaches that predict protein-protein interactions, including Direct Coupling Analysis (DCA), represent an attractive alternative to dissect interactions within protein complexes. However, due to high computational complexity and high false positive rate they are applicable only to small proteins. Here, we present a modified DCA to predict residues and domains involved in interactions of large proteins. To reduce false positive levels and increase accuracy of prediction, we use local Gaussian averaging and predicted secondary structure elements. As a proof-of-concept, we apply our method to two Integrator subunits, INTS9 and INTS11, which form a heterodimeric structure previously solved by crystallography. We accurately predict the domains of INTS9/11 interaction. We then apply this approach to predict the interaction domains of two complexes whose structure is currently unknown: 1) The heterodimer formed by the Cleavage and Polyadenylation Specificity Factor 100-kD (CPSF100) and 73-kD (CPSF73); 2) The heterotrimer formed by INTS4/9/11. Our predictions of interactions within these two complexes are supported by experimental data, demonstrating that our modified DCA is a useful method for predicting interactions and can easily be applied to other complexes.

## INTRODUCTION

Traditional methods to study protein association, including the yeast two-hybrid and co-immunoprecipitation analyses, have been shown to be reliable in characterizing protein complexes but they remain laborious and time consuming. Therefore, computational methods to predict protein-protein interactions are an attractive alternative to experimental methods. One of such methods is evolutionary coupling analysis. The underlying idea of evolutionary coupling is that, to preserve function, a mutation in one of interacting residues is likely to be compensated by a complementary mutation in the other. The key advantage of this approach is that interactions between residues are detected not only based on their physical proximity (as in co-crystallization studies), but on evolutionary pressure, and therefore more likely functional. Co-evolution of residues in protein sequences has been used for predicting residue-residue interactions (contacts) in small, bacterial proteins for a few decades (Lockless and Ranganathan 1999; Morcos et al. 2011; Ochoa et al. 2013). As observed by us and others (Hopf et al. 2012; Hopf et al. 2014b; Fongang et al. 2019), with the rapid increase in number of sequenced animal genomes, these methods also became feasible for protein interactions in metazoans, including humans, and for larger proteins and even protein complexes (Hopf et al. 2012; Naveed et al. 2012; Hopf et al. 2014b; Kim et al. 2014; Ovchinnikov et al. 2014; dos Santos et al. 2015; Tesileanu et al. 2015; Abriata et al. 2016; Champeimont et al. 2016; Feinauer et al. 2016; Lua et al. 2016; Neuwald 2016; Yu et al. 2016).

Specific interactions between proteins impose evolutionary constraints on the interacting partners. For instance, mutation of a contact residue in one partner generally impairs binding but may be compensated by a complementary mutation in the other partner. Such coevolution of interaction partners results in correlations between their amino acid sequences than can be observed by analyzing Multiple Sequence Alignment (MSA) of the interacting proteins across multiple species, and can be used to predict residue– residue contacts (Marks et al. 2011; Morcos et al. 2011; Morcos et al. 2014; Wang et al. 2017). Recently, methods from statistics and statistical physics were used to disentangle direct couplings, related to actual residue interactions, from mere correlations between MSA columns. This has resulted in a new class of methods, including the Direct Coupling Analysis (DCA), that can reliably predict protein structures using only sequence information provided enough homologous sequences are available. However, the applications of DCA methods to large proteins are limited by high number of typically generated false positives. To reduce the number of false positives we have recently proposed to use a local convolution of evolutionary coupling (EC) scores with a kernel fitted to the structure of the proteins (Fongang et al. 2019). Here, we will adapt DCA to study interactions of the Integrator complex subunits, which are particularly challenging due to size of some of the subunits.

The Integrator complex (INT) is an important transcriptional component in regulating 3’-end processing of non-coding RNA (Baillat and Wagner 2015), and widely participating in transcription processes at protein-coding genes through its association with RNA polymerase II (Gardini et al. 2014; Stadelmayer et al. 2014). Several INT subunits have been found to play important roles in human brain development(Oegema et al. 2017), lung function(Kheirallah et al. 2017), embryogenesis(Kapp et al. 2013), and adipose differentiation(Otani et al. 2013). Among at least the 14 subunits of Integrator, Subunits 9 and 11 (INTS9-INTS11) constitute a catalytic core of INT and are paralogs of two 100 kDa and 73 kDa subunits of the Cleavage and Polyadenylation Specificity Factor (CPSF) (Wu et al. 2017; Albrecht et al. 2018). INTS11 forms a stable complex with INTS9 through their C-terminal domains (CTDs) that also exists in CPSF73 and CPSF100 but with poor sequence conservation(Albrecht and Wagner 2012; Albrecht et al. 2018). Recently, the crystal structure of the INTS9/INTS11 CTD complex has been reported at 2.1 Å resolution, which explains high binding affinity for the two proteins(Wu et al. 2017). Moreover, the binding of INTS9/INTS11 has been shown to be a prerequisite to the recruitment of INTS4 which altogether form the INTS4/9/11 heterotrimer (Albrecht et al. 2018), believed to be the most critical subunits for snRNA 3′-end formation. Like their INT counterparts, CPSF73 and CPSF100 are present in a stable complex and are also required for 3’-end processing of all metazoan pre-mRNAs (Sullivan et al. 2009). Although the crystal structures of human CPSF73 and yeast CPSF100(Mandel et al. 2006) have been reported, the molecular basis of their interaction as well the mechanism leading to the CPSF73/100/SYM trimerization are still unknown.

Here, we use DCA with our added convolution method to accurately predict the binding residues of INTS9/INTS11 heterodimer. We also study the interaction between INTS9 and INTS11 paralogs, CPSF73 and CPSF100, and identify their most likely binding interfaces to be the C-terminal domains of both proteins. Since it has been suggested that INTS4, INTS9, and INTS11 form a heterotrimer, we used our coevolution method to show that such trimerization involves the N-, and C-terminal domains of INTS4 exclusively. As the structure of the Integrator Complex remains largely unknown, experimental validation of these predictions will set the foundation for the complete description of the structure of the whole complex.

## RESULTS

### Modified Direct Coupling Analysis (DCA) approach

DCA algorithms have been shown to produce a large number of false positives, but we recently suggested (Fongang et al. 2019) that post-processing of DCA map data, based on the local convolution with Gaussian kernel, may lead to reducing noise in the prediction of most likely interacting residues (Fongang et al. 2019). A schematic description of the method is presented in Fig. 1. To reduce running time, we used a variant of DCA, the pseudo-likelihood maximization Direct-Coupling Analysis (*plmDCA*) (Ekeberg et al. 2014), which has a lower computational cost than traditional DCA. Evolutionary coupling score (ECs) was calculated and used to build the corresponding coupling matrix.

**Figure 1.**
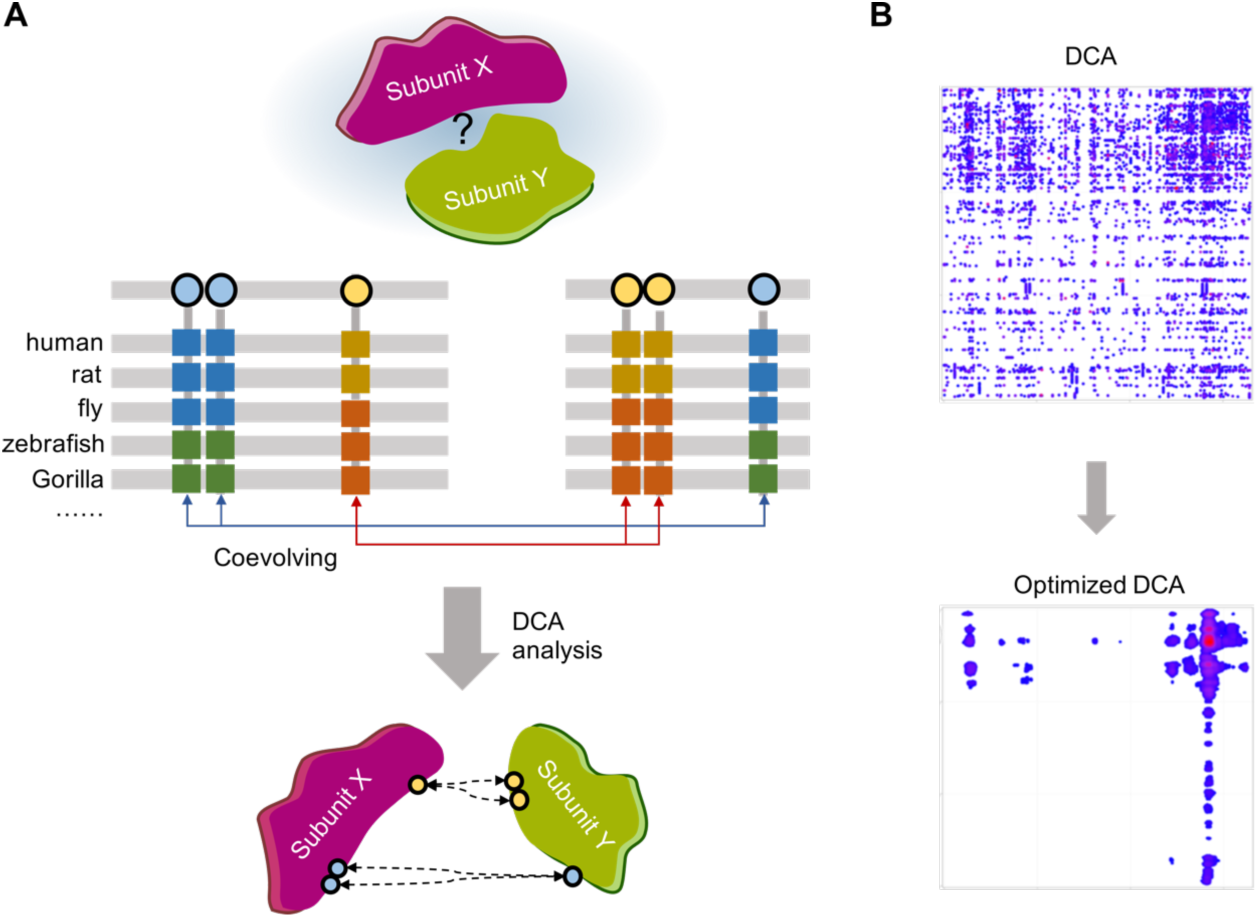
Prediction of protein-protein interactions based on co-evolutionary analysis. (a) The principle of co-evolution analysis. Coevolution between residues of interacting proteins can be used to predict the binding interfaces as described by Hopf et al. (Hopf et al. 2014a). (b) Optimizing the coevolution maps. For large proteins the predictions are hindered by false positives resulting from statistical background noise. Genuine interactions between proteins generally involve stretches of residues rather than individual amino acids, therefore local convolution of evolutionary scores and structural properties can reduce the noise and filter out the false positives, allowing to correctly identify the interacting regions, as described by Fongang et al. (Fongang et al. 2019).

To improve our prediction and avoid false positive in DCA analysis, we use local convolution of ECs, derived from our approach introduced previously in (Fongang et al. 2019). Gaussian convolution is applied to local structural elements (here defined by secondary structure) to count the contribution of neighboring residues with an assumption that contacts between proteins occur locally and drive residues evolving within the same structural elements. In our experience, an isolated strong EC peak surrounded by low EC values for residues belonging to different secondary structure elements is more likely to be a false positive than a cluster of less high EC values for residues in the same secondary structure element. Therefore, a convolution of EC scores with a kernel based on secondary structure information can predict the more likely interacting residues. The convolution algorithm depends on several parameters including the length of the interacting residues, the variances of the Gaussian and the predicted structures. The convolved EC score for a pair of residues (*i,j*) is

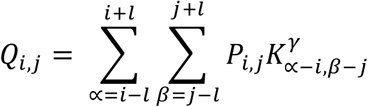

where

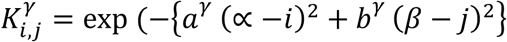

is the kernel function fitted on the structural elements and *P*_*i,j*_ are the EC scores computed using the evolutionary coupling algorithm. The key innovation over the previous variant of the method we proposed is to derive the values of the parameters *a*^*γ*^, *b*^*γ*^, based on the details of interaction as characterized by a previous crystallographic study. Second, based on previous experience, and on the argument that an interaction interface is expected to affect both proteins in a statistically similar manner, we decided to only consider convolution models assuming *a*^*γ*^ *= b*^*γ*^ for every value of *γ*. These parameters were optimized using the INTS9/INTS11 complex for which structural information of their interacting C-terminal domains was available. We started the optimization from Evolutionary Coupling (EC) scores of INTS9/INTS11, which were generated based on multiple sequence alignment of both proteins over 204 species. Next, we converted EC scores into DCA maps representing interactions between residues of the two proteins (Fig. 2a). The optimal values of *a*^*γ*^, *b*^*γ*^, and *l* were selected to maximum overlap between the prediction and the experimental distances for INTS9/11 interaction and used also for other cases for which crystal structures are unknown. The optimized parameters for the Gaussian convolution are provided in Table 3.

**Table 1:** Characterization of the different complexes studied. The global p-value was computed using the simulated phylogeny as described by Fongang et al. (Fongang et al. 2019). Briefly, we randomized the sequence distribution of one protein while keeping the other unchanged and computed the ECs of the interaction. The p-value was then estimated by counting the ECs higher than a predefined cutoff for the 100 randomizations. (*** denotes the most likely and * second most likely binding region, N-C means N-terminal domain of protein A and C-terminal domain of protein B. When necessary, the specific amino-acids are indicated).

| **Protein A** | **Protein B** | **Sequences in cMSA** | **Global p-value for interaction** | **Most likely interacting domains** |
| --- | --- | --- | --- | --- |
| INTS9 | INTS11 | 204 | <0.01 | C-C (***), N-C (*) |
| INTS4 | INTS9 | 171 | 0.01 | N-N, N-C, C-N, or C-C |
| INTS4 | INTS11 | 171 | 0.01 | N-N, N-C, C-N, or C-C |
| CPSF100 | CPSF73 | 138 | 0.04 | C-C (***), C-R344-525(*) |

**Table 2:** Properties of the protein heterodimers studied. The significant length (number of sequences in cMSA / Size of both proteins) is indicated for each protein complex. As they are all lower than the required 0.7 for most DCA analysis in single proteins (Hopf et al. 2014a), we used our method, based on local convolution of ECs, to reduce the number of false positive generated and highlight the most likely interacting partners (Fongang et al. 2019).

|  | Protein size (AA) | Number of individual sequences | Number of sequences in cMSA | Sig. length | Sequence coverage |
| --- | --- | --- | --- | --- | --- |
| INTS9 | 658 | 223 | 204 | 0.16 | 55 – 90% |
| INTS11 | 600 | 239 |  |  |  |
| INTS4 | 963 | 202 | 171 | 0.11 | 55 – 90% |
| INTS9 | 658 | 223 |  |  |  |
| INTS4 | 963 | 202 | 171 | 0.11 | 55 – 90% |
| INTS11 | 600 | 239 |  |  |  |
| CPSF100 | 782 | 179 | 138 | 0.1 | 55 – 90% |
| CPSF73 | 684 | 161 |  |  |  |

**Table 3:** Optimized values of the Gaussian convolution. *a* and *b* are the optimized variances of the Gaussian Kernel, *l* is the average length of the interval with interacting residues and γ = 1 if the residues belong to the same secondary structure, if not γ = 2.

| | $a$ | $b$ | $l$ (AA) |
| --- | --- | --- | --- |
| $\gamma = 1$ | 0.01 – 0.05 | 0.01 – 0.05 | 17 - 24 |
| $\gamma = 2$ | 0.001 - 0.008 | 0.001 – 0.008 | 17 - 24 |

**Figure 2.**
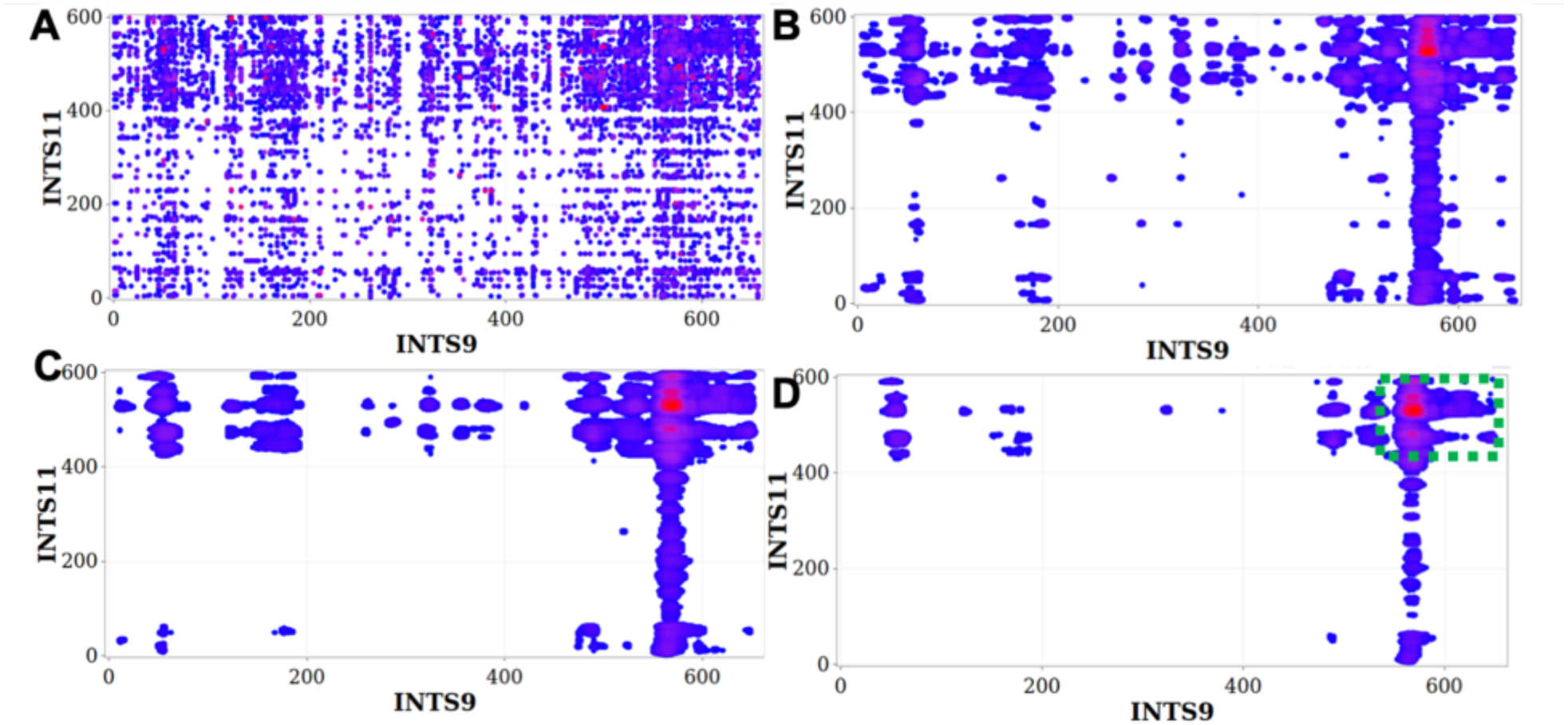
Convolved ECs of INTS9/INTS11 reveal most likely interacting residues. The local convolution method was applied to the top 1% of raw ECs (A) using different parameters representing the length of the interacting residues *l*, and the variances of the Gaussian kernels *a* and *b*. (B): *l* = 18 AA, *a* = *b* = 0.1; (C): *l* = 21AA, *a* = *b* = 0.05; (D): *l* = 21AA, *a* = *b* = 0.01. The green rectangle on (D) delimits the most likely interacting region.

### Modified DCA correctly predicts the interacting residues of INTS9/INTS11 heterodimer

To validate our method we applied it to predict interactions of the INTS9 and INTS11 heterodimer, which interface has been solved by crystallography. Raw DCA method applied to predicting INTS9/11 interactions generated a very high level of statistical background noise in the EC map leading to false positives thereby making identification of binding residues very challenging (Fig. 2a). Therefore, we applied our local convolution algorithm (Fongang et al. 2019) to the EC map of INTS9/INTS11 with variable parameters describing the variances of the Gaussian kernel and the lengths of stretches of interacting residues (Fig. 2b-2d). Indeed, as we changed the convolution parameters, stretches of residues starting at residue 553 of INTS9 and 422 of INTS11 yielded highly optimized co-evolutionary scores compared to the entire DCA map. The optimized parameters corresponding to the final convolved DCA map (Fig. 2D) are l (interacting residues) = 21 AA, a=b (Gaussian variances) = 0.01. The result of applying this algorithm indicated that the C-terminal domains of both INTS9 and INTS11 contain the most likely interacting residues of the INTS9/INTS11 heterodimer.

Experimental evidence demonstrates that INTS9 and INTS11 interact through their C-terminal domains (CTD). Indeed, the report by Wu et al. (Wu et al. 2017) explains the molecular basis and the functionality of the INTS9/INTS11 heterodimer as well as the crystal structure of the CTD at 2.1 Å. To assess the accuracy of our predictions, we compared the pairs of residues (one from INTS9, one from INTS11) at distances less than 6.0 Å to the pairs of interacting residues we predicted. Using this criterion for comparison, we observe very good agreement between our modified DCA predictions and the crystal structure of the IntS9/11 CTD. We find that 73% of the pairs with top 5% highest convolved signals are within the CTD of both proteins and 81% of the experimentally determined contacts were predicted by our method. Also, convolved ECs are correlated with structural information including solvent accessibility, secondary structure, and physical-chemical properties. This test show that our method has potential to accurately predict residues involved in protein-protein interactions. It also confirms that the previously observed contact between the CTDs of INTS9 and INTS11 is a physiologically significant interaction that is subject to positive evolutionary pressure. Finally, we need to note that we use the INTS9/INTS11 structure as validation of the method while we also used the same interaction for optimizing the parameters for the convolution procedure. Nonetheless, the amount of information recycled here is minimal – the INTS9/11 structure was only used to optimize the values of the three parameters (*a*^*γ*^, *b*^*γ*^, and *l*), while the validation is based on a very large number (>10^5^) of EC scores.

### Predicting Interacting interfaces of the CPSF100/CPSF73 heterodimer

The validation of the predicted INTS9/INTS11 interfaces allowed us to expand our study to the Cleavage and Polyadenylation Specificity Factor (CPSF) complex which is involved in the 3’-end cleavage of pre-mRNA prior to polyadenylation. Within the CPSF complex, CPSF73 has been shown to form a stable and functional heterodimer with CPSF100 but the molecular basis for this interaction is not known. Moreover, CPSF100 and CPSF73 are structurally very similar to INTS9 and INTS11 and are annotated as paralogs, respectively. INTS11 contains its highest degree of conservation with CPSF73 over much of the N-terminal MBL domains and the β-CASP domains but are highly divergent at the C-terminal regions (Albrecht and Wagner 2012). CPSF100 and INTS9 are both inactivated through changes in key catalytic residues, but INTS9 with a molecular mass of 74kDa is much smaller than CPSF100. Although both sets of proteins are similar and the INTS9/INTS11 structure has been solved, little is known about the structural basis of the CPSF100/CPSF73 heterodimer.

As with INTS9/INTS11, we used 138 pairs of CPSF100 and CPSF73 orthologous sequences from metazoans to compute DCA maps of both proteins (Fig. 4a). Then we applied the local convolution of ECs scores with optimized parameters obtained from the previous case to highlight the most likely interacting residues (Fig. 4b-d). This analysis predicts that the most likely interacting residues of CPSF100/CPSF73 involve their respective C-terminal domains (Fig. 4d, Fig. 5). Indeed 88% of the top 5% highest convolved EC scores are within the region comprising the last 115 and 164 amino acids of CPSF100 and CPSF73, respectively. The second most likely interacting residues comprised the region from 367-533 on CPSF100 and the CTD of CPSF73 (Fig. 4d). This information is similar to INTS9/INTS11 and confirmed the striking similarity between the two complexes. Moreover, the findings obtained using our modified DCA method are consistent with those obtained biochemically by Michalski *et al*. (Michalski and Steiniger 2015) in that both the C-terminal domains of CPSF100 and CPSF73 are required for the core cleavage complex formation.

**Figure 3.**
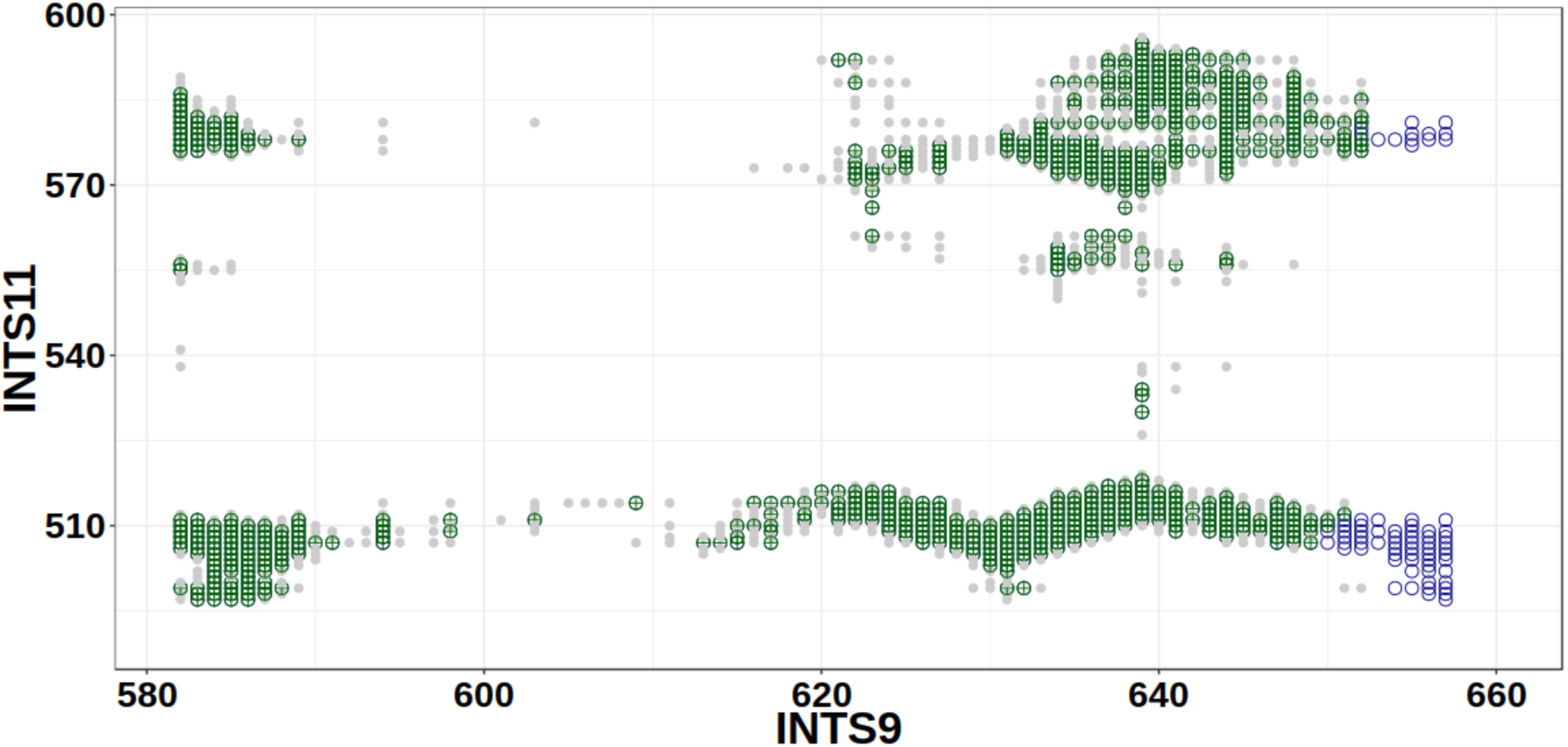
Example of quality of contact inference. After refinement by EC averaging and secondary structure information, our contact predictions are highly consistent with crystallographic contacts from the recent structure of the INTS9/INTS11 dimer. Green: predicted contacts that were confirmed experimentally, gray: experimental contacts that were not predicted, blue empty circles: predicted contacts that were not experimentally confirmed. X- and Y-axes: positions in INTS9 and INTS11 (only the interacting C-terminal region is shown).

**Figure 4.**
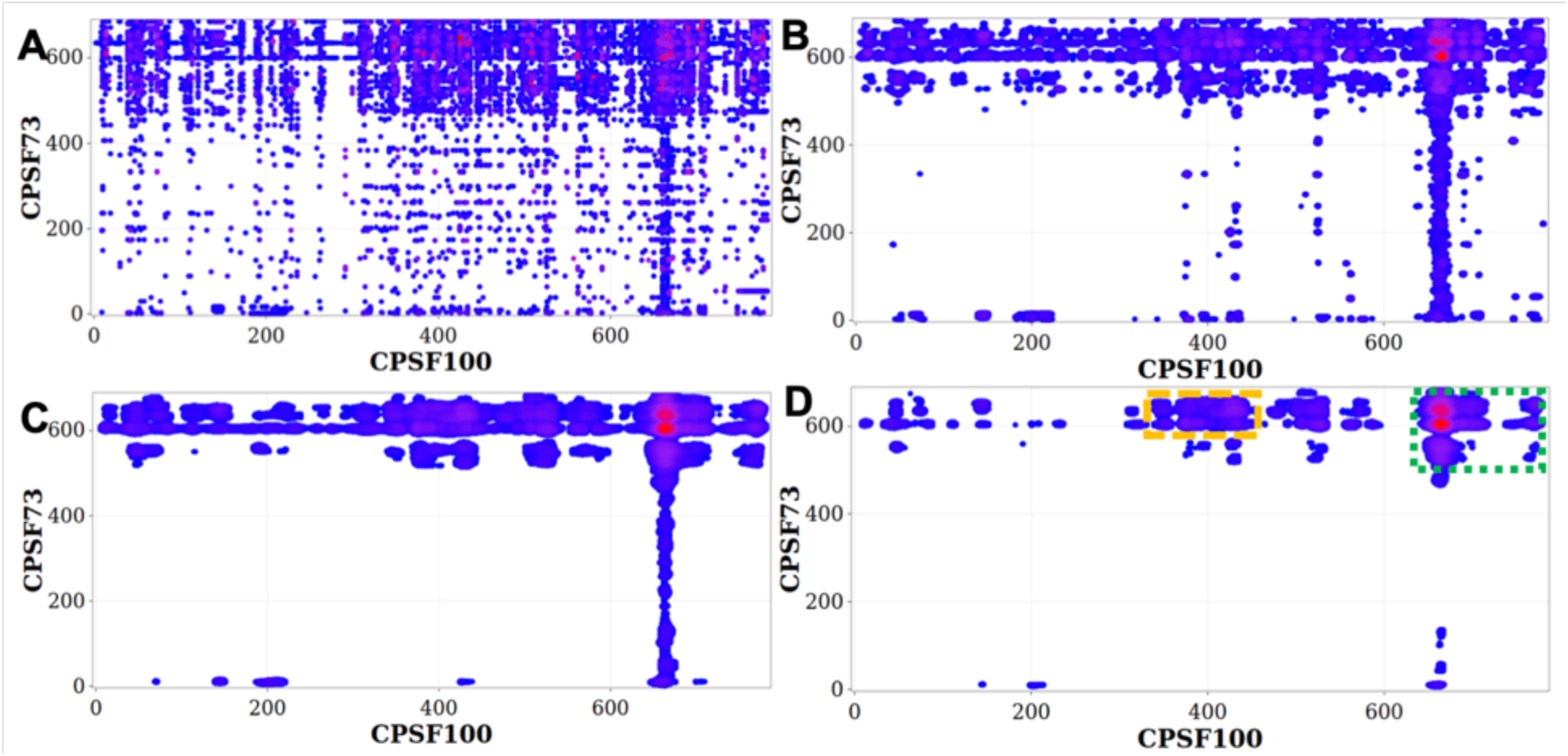
Convolved ECs of CPSF73 – CPSF100 reveal most likely interacting residues. The local convolution method was applied to the top 1% of raw ECs (A) using different parameters representing the length of the interacting residues *l*, and the variances of the Gaussian kernels *a* and *b*. (B): *l* = 18 AA, *a* = *b* = 0.1; (C): *l* = 21AA, *a* = *b* = 0.05; (D): *l* = 21AA, *a* = *b* = 0.01. The green and orange rectangles on (D) delimit the most likely and the second likely interacting regions of the CPSF100/CPSF73 heterodimer respectively.

**Figure 5.**
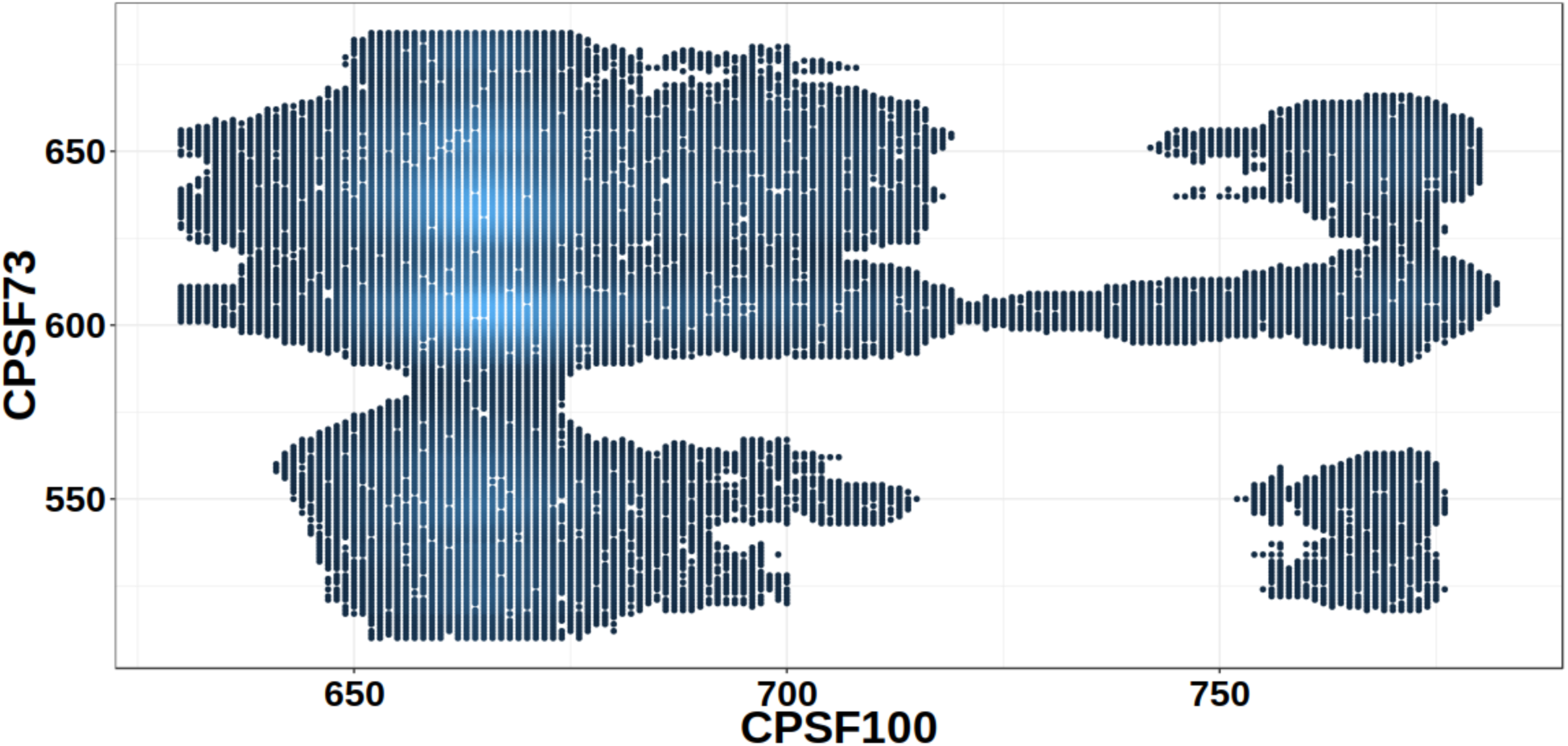
Optimized coevolution map of the C-Terminal Domains from the CPSF100/CPSF73 heterodimer. The figure shows the predicted and optimized EC scores (*l* = 21AA, a = b 0.01, top 1%). Grey indicates the predicted interactions, blue-most likely interactions.

### Predicting interacting residues of the INTS4/9/11 heterotrimer

Encouraged by results of our modified DCA approach for the above two heterodimers, we applied our method to a heterotrimeric complex. Predicting structure of heterotrimer presents an additional challenge to current DCA analyses because in a heterotrimer also indirect interactions may present as couplings between residues, thus significantly increasing the number of possible indirect links between the residues (interaction between a residue Ai in subunit A with any of the hundreds of residues in subunit B, combined with interaction between the residues in B and residue Cj in subunit C may be interpreted as an interaction between Ai and Cj). This effect may increase the number of false positives that may affect the results of the analysis. Reducing the number of artifacts through post-processing of the EC map we proposed is therefore even more important in the case of predicting a contact interface between two subunits within a heterotrimeric complex. INTS4 has been reported to associate with INTS9 and INTS11 to form the INTS4/9/11 heterotrimer and has been proposed to behave similar to how Symplekin associates with CPSF100/73. Importantly, the molecular basis of both heterotrimers has not been established. Using the *plmDCA* algorithm coupled to the convolution of resulting ECs as described in the Methods section, we predicted that the N- and C-terminal domains of INTS4 are both interacting with INTS9/11 (Fig. 6). Our results also suggests that INTS4 has ability to bind both INTS9 and INTS11 at the same time, but coevolutionary analysis was unable to significantly distinguish which of the N- or C-terminal was associated with INTS9 (or INTS11). Interestingly, this model is strikingly similar to the heterotrimerization of CPSF100/CPSF73/Symplekin as proposed by Michalski et al. (Michalski and Steiniger 2015) and is consistent with previously published biochemical experiments analyzing binding domains involved in the INTS4/9/11 heterotrimer.

**Figure 6.**
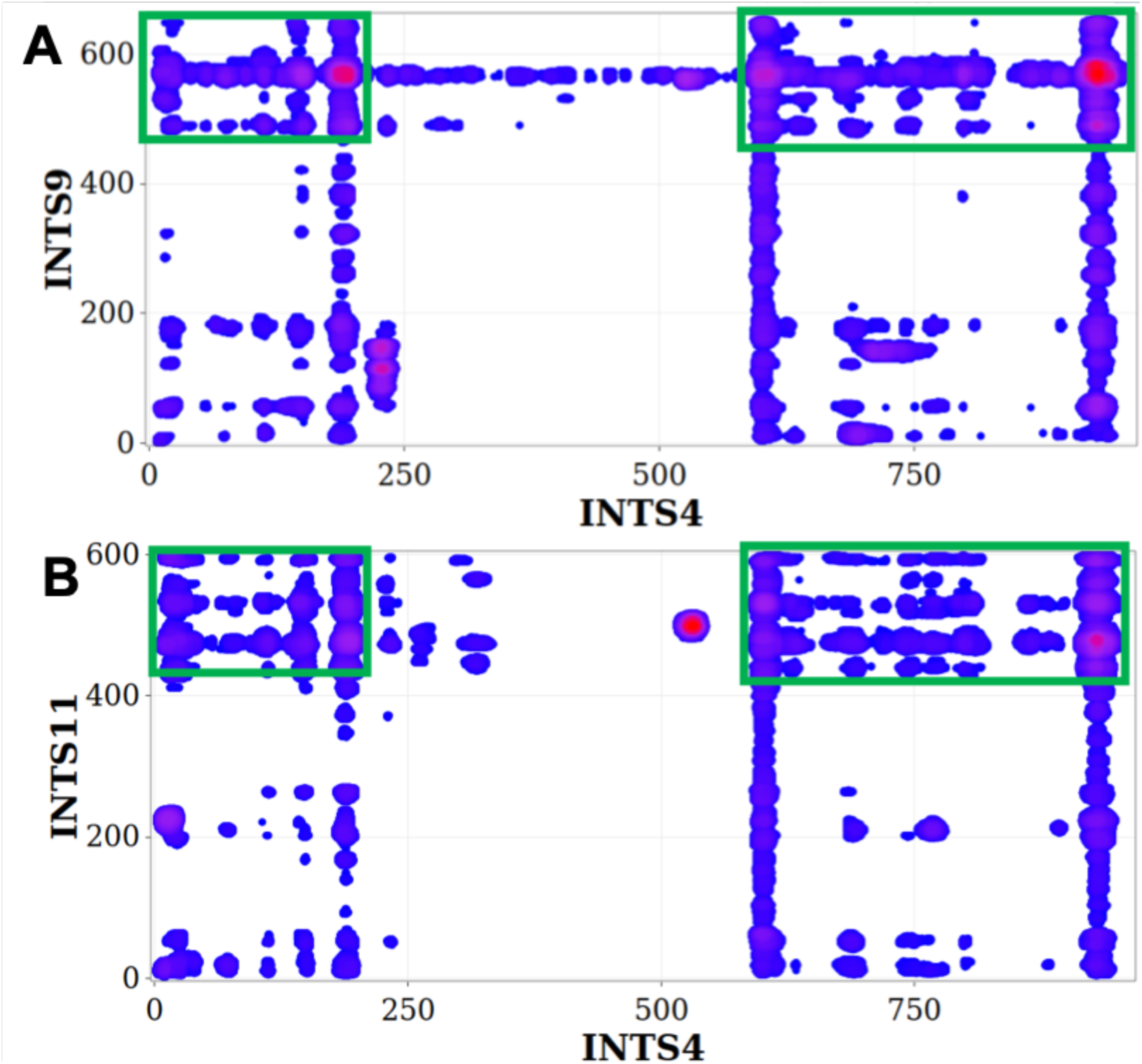
The predicted contacts between INTS4/INTS9 and INTS4/INTS11 involve the C- and N-terminal domains of INTS4. The figure shows the convolved ECs (*l* = 21AA, *a* = *b* = 0.01, top 1%) of the INTS4/INTS9 (A) and INTS4/INTS11 (B).

**Figure 7.**
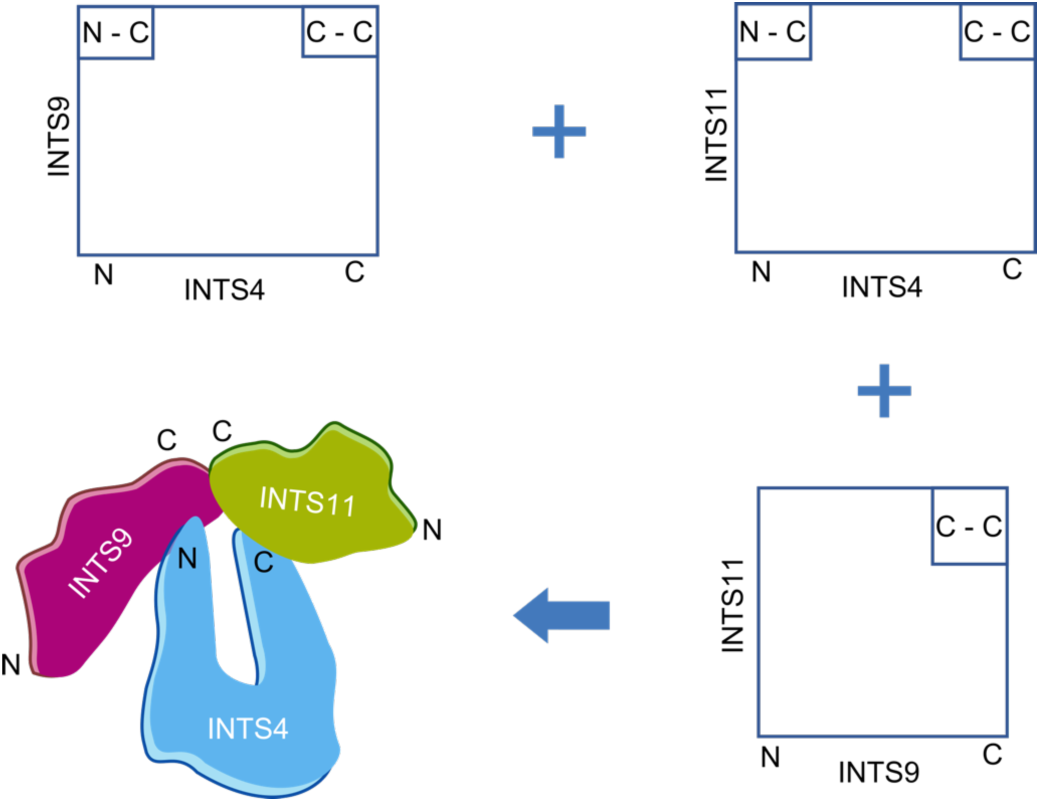
Schematic view of the predicted structure of INTS4/9/11 trimer. Both the N- and C-terminal domains of INTS4 are involved in the interaction with C-terminal domains of INTS9 and INTS11.

## DISCUSSION

In this study, we applied coevolutionary method to identify the most likely binding residues of INTS4, INTS9, and INTS11 as well as CPSF73 and CPSF100. We modified the Direct Coupling Analysis (DCA) algorithm with several changes we introduced to allow more accurate inference of interactions. Specifically, we used local Gaussian convolution and predicted secondary structure to reduce number of false positives and thus increase the accuracy of predicted interactions. As discussed above, coevolution between residues of interacting proteins can be used to predict the binding interfaces. However, for large proteins, the predictions are hindered by statistical background noise and the large number of false positives that are generated. Because an interaction between proteins generally involves stretches of residues rather than individual amino acids, in our experience local convolution of evolutionary scores and structural properties tend to predict more accurately the most likely interacting residues (Fongang et al. 2019).

As proof-of-concept we inferred the binding interface of INTS9/INST11 heterodimer using our local convolution algorithm (Fongang et al. 2019). Our results show that the most likely interacting residues of the INTS9/INTS11 heterodimer are located on the C-terminal domains of both INTS9 and INTS11. This predicted result has been validated by Wu *et al*. (Wu et al. 2017) through the U7-GFP reporter, by which they found that the mutations specifically disrupt the formation of the INTS9/INTS11 C-terminal domain heterodimer. Thus, our method has been validated to be accurate in predicting the binding residues of large proteins. Then, we applied our method to another two important proteins in the cleavage of pre-mRNA: CPSF100 and CPSF73, which are paralogs of INTS9 and INTS11, respectively. However, their interactions are unknown. Our prediction confirmed that both the C-terminal domains of CPSF100 and CPSF73 are required for the core cleavage complex formation *in vivo* and the binding with Symplekin as reported by Michalski *et al*. (Michalski and Steiniger 2015). We found that the N- and C-terminal domains of INTS4 could interact with INTS9 and INTS11, as shown in Fig. 6, suggesting INTS4 has ability to bind both INTS9 and INTS11.

Using coevolution of residues, we have characterized several interactions between proteins related to the Integrator complex. The results are summarized in Table 2. Our analysis confirms the physiological nature of several interactions indicated by previous studies, and predicts new interactions that can help to explain the nature of the Integrator, a complex molecular machine. Finally, the predicted characteristics of the interactions between pairs of proteins and the identification of domains and residues potentially crucial for the respective dimerization can inform future experimental studies, such as targeted mutations that may disrupt complex formation.

## MATERIALS AND METHODS

### Protein sequence collection and alignment

The initial step in studying evolutionary coupling between proteins is to construct a concatenated Multiple Sequences Alignment (cMSA) by aligning orthologs of proteins under study and join the alignment by species. It is important than both orthologs come from the same species. INTS4, INTS9, INTS11, CPSF73, and CPSF100 have all been shown to be well conserved across metazoans, with sequences comprising the β-Lactamase, β-CASP and C-terminal domains. Their sequences were extracted by querying GENBANK (Benson et al. 1993) and by running genome-wide *tblastn* (Altschul et al. 1990) against genomes not present in GENBANK. The sequences were aligned to human reference sequence, and those with 60-90% similarity were used. 223, 239, 202, 179, and 161 orthologs of INTS9, INTS11, INTS4, CPSF100, and CPSF73 respectively were obtained. Common orthologs for a pair of proteins were aligned using Clustal-Ω (Sievers et al. 2011) (see Table 1).

### Prediction of protein heterodimers and heterotrimers

Evolutionary couplings between INTS9/INTS11, INTS4/INTS9, INTS4/INTS11 and CPSF100/CPSF73 were analyzed using Direct Coupling Analysis (DCA) algorithm (Morcos et al. 2011), which can distinguish direct functional residue interactions from correlations resulting from indirect interactions and assign higher scores to direct correlations rather than to indirect ones. To reduce running time, we used a variant of DCA, the pseudo-likelihood maximization Direct-Coupling Analysis (*plmDCA*) (Ekeberg et al. 2014), which has a lower computational cost than traditional DCA. Evolutionary coupling scores (ECs) were calculated and used to build the corresponding coupling matrix. To improve our prediction and avoid false positives in DCA analysis, we used local convolution of ECs as described and validated above.

## ACKNOWLEDGMENTS

The study was supported in part by the NIH GM grant R01 GM112131, a training fellowship from the Gulf Coast Consortia Computational Cancer Biology Training Program (CPRIT Grant No. RP170593) awarded to Y.Z, by a pilot grant from the Center for Addiction Research at UTMB and by The Welch Foundation [H1889 to E.J.W.].

## REFERENCES

Abriata LA, Bovigny C, Dal Peraro M. 2016. Detection and sequence/structure mapping of biophysical constraints to protein variation in saturated mutational libraries and protein sequence alignments with a dedicated server(vol 1, 242, 2016). Bmc Bioinformatics 17.

Albrecht TR, Shevtsov SP, Wu Y, Mascibroda LG, Peart NJ, Huang KL, Sawyer IA, Tong L, Dundr M, Wagner EJ. 2018. Integrator subunit 4 is a ‘Symplekin-like’ scaffold that associates with INTS9/11 to form the Integrator cleavage module. Nucleic acids research 46: 4241–4255.

Albrecht TR, Wagner EJ. 2012. snRNA 3’ end formation requires heterodimeric association of integrator subunits. Molecular and cellular biology 32: 1112–1123.

Altschul SF, Gish W, Miller W, Myers EW, Lipman DJ. 1990. Basic local alignment search tool. J Mol Biol 215: 403–410.

Baillat D, Wagner EJ. 2015. Integrator: surprisingly diverse functions in gene expression. Trends in biochemical sciences 40: 257–264.

Benson D, Lipman DJ, Ostell J. 1993. GenBank. Nucleic Acids Res 21: 2963–2965.

Champeimont R, Laine E, Hu SW, Penin F, Carbone A. 2016. Coevolution analysis of Hepatitis C virus genome to identify the structural and functional dependency network of viral proteins. Scientific Reports 6.

Chen J, Waltenspiel B, Warren WD, Wagner EJ. 2013. Functional analysis of the integrator subunit 12 identifies a microdomain that mediates activation of the Drosophila integrator complex. The Journal of biological chemistry 288: 4867–4877.

dos Santos RN, Morcos F, Jana B, Andricopulo AD, Onuchic JN. 2015. Dimeric interactions and complex formation using direct coevolutionary couplings. Scientific Reports 5.

Ekeberg M, Hartonen T, Aurell E. 2014. Fast pseudolikelihood maximization for direct-coupling analysis of protein structure from many homologous amino-acid sequences. Journal of Computational Physics 276: 341–356.

Elking DM, Fusti-Molnar L, Nichols A. 2016. Crystal structure prediction of rigid molecules. Acta Crystallogr B Struct Sci Cryst Eng Mater 72: 488–501.

Feinauer C, Szurmant H, Weigt M, Pagnani A. 2016. Inter-Protein Sequence CoEvolution Predicts Known Physical Interactions in Bacterial Ribosomes and the Trp Operon. Plos One 11.

Fongang B, Cunningham KA, Rowicka M, Kudlicki A. 2019. Coevolution of Residues Provides Evidence of a Functional Heterodimer of 5-HT2AR and 5-HT2CR Involving both Intracellular and Extracellular Domains. Neuroscience doi: https://doi.org/10.1016/j.neuroscience.2019.1005.1013.

Gardini A, Baillat D, Cesaroni M, Hu D, Marinis JM, Wagner EJ, Lazar MA, Shilatifard A, Shiekhattar R. 2014. Integrator regulates transcriptional initiation and pause release following activation. Molecular cell 56: 128–139.

Hopf TA, Colwell LJ, Sheridan R, Rost B, Sander C, Marks DS. 2012. Three-Dimensional Structures of Membrane Proteins from Genomic Sequencing. Cell 149: 1607–1621.

Hopf TA, Scharfe CP, Rodrigues JP, Green AG, Kohlbacher O, Sander C, Bonvin AM, Marks DS. 2014a. Sequence co-evolution gives 3D contacts and structures of protein complexes. Elife 3.

Hopf TA, Scharfe CPI, Rodrigues JPGLM, Green AG, Kohlbacher O, Sander C, Bonvin AMJJ, Marks DS. 2014b. Sequence co-evolution gives 3D contacts and structures of protein complexes. Elife 3.

Kapp LD, Abrams EW, Marlow FL, Mullins MC. 2013. The integrator complex subunit 6 (Ints6) confines the dorsal organizer in vertebrate embryogenesis. PLoS genetics 9: e1003822.

Kheirallah AK, de Moor CH, Faiz A, Sayers I, Hall IP. 2017. Lung function associated gene Integrator Complex subunit 12 regulates protein synthesis pathways. BMC genomics 18: 248.

Kim DE, DiMaio F, Wang RYR, Song YF, Baker D. 2014. One contact for every twelve residues allows robust and accurate topology-level protein structure modeling. Proteins-Structure Function and Bioinformatics 82: 208–218.

Lockless SW, Ranganathan R. 1999. Evolutionarily conserved pathways of energetic connectivity in protein families. Science 286: 295–299.

Lua RC, Wilson SJ, Konecki DM, Wilkins AD, Venner E, Morgan DH, Lichtarge O. 2016. UET: a database of evolutionarily-predicted functional determinants of protein sequences that cluster as functional sites in protein structures. Nucleic Acids Research 44: D308–D312.

Mandel CR, Kaneko S, Zhang H, Gebauer D, Vethantham V, Manley JL, Tong L. 2006. Polyadenylation factor CPSF-73 is the pre-mRNA 3’-end-processing endonuclease. Nature 444: 953–956.

Marks DS, Colwell LJ, Sheridan R, Hopf TA, Pagnani A, Zecchina R, Sander C. 2011. Protein 3D structure computed from evolutionary sequence variation. PLoS One 6: e28766.

Marks DS, Hopf TA, Sander C. 2012. Protein structure prediction from sequence variation. Nature Biotechnology 30: 1072–1080.

Michalski D, Steiniger M. 2015. In vivo characterization of the Drosophila mRNA 3’ end processing core cleavage complex. RNA 21: 1404–1418.

Morcos F, Hwa T, Onuchic JN, Weigt M. 2014. Direct coupling analysis for protein contact prediction. Methods Mol Biol 1137: 55–70.

Morcos F, Pagnani A, Lunt B, Bertolino A, Marks DS, Sander C, Zecchina R, Onuchic JN, Hwa T, Weigt M. 2011. Direct-coupling analysis of residue coevolution captures native contacts across many protein families. Proceedings of the National Academy of Sciences of the United States of America 108: E1293–E1301.

Naveed H, Xu Y, Jackups R, Liang J. 2012. Predicting Three-Dimensional Structures of Transmembrane Domains of beta-Barrel Membrane Proteins. Journal of the American Chemical Society 134: 1775–1781.

Navio D, Rosell M, Aguirre J, de la Cruz X, Fernandez-Recio J. 2019. Structural and Computational Characterization of Disease-Related Mutations Involved in Protein-Protein Interfaces. Int J Mol Sci 20.

Neuwald AF. 2016. Gleaning structural and functional information from correlations in protein multiple sequence alignments. Current Opinion in Structural Biology 38: 1–8.

Ochoa D, Garcia-Gutierrez P, Juan D, Valencia A, Pazos F. 2013. Incorporating information on predicted solvent accessibility to the co-evolution-based study of protein interactions. Molecular Biosystems 9: 70–76.

Oegema R, Baillat D, Schot R, van Unen LM, Brooks A, Kia SK, Hoogeboom AJM, Xia Z, Li W, Cesaroni M et al. 2017. Human mutations in integrator complex subunits link transcriptome integrity to brain development. PLoS genetics 13: e1006809.

Otani Y, Nakatsu Y, Sakoda H, Fukushima T, Fujishiro M, Kushiyama A, Okubo H, Tsuchiya Y, Ohno H, Takahashi S et al. 2013. Integrator complex plays an essential role in adipose differentiation. Biochemical and biophysical research communications 434: 197–202.

Ovchinnikov S, Kamisetty H, Baker D. 2014. Robust and accurate prediction of residueresidue interactions across protein interfaces using evolutionary information. Elife 3.

Pearce R, Huang X, Setiawan D, Zhang Y. 2019. EvoDesign: Designing Protein-Protein Binding Interactions Using Evolutionary Interface Profiles in Conjunction with an Optimized Physical Energy Function. J Mol Biol.

Price SL. 2004. The computational prediction of pharmaceutical crystal structures and polymorphism. Adv Drug Deliv Rev 56: 301–319.

Sievers F, Wilm A, Dineen D, Gibson TJ, Karplus K, Li WZ, Lopez R, McWilliam H, Remmert M, Soding J et al. 2011. Fast, scalable generation of high-quality protein multiple sequence alignments using Clustal Omega. Mol Syst Biol 7.

Stadelmayer B, Micas G, Gamot A, Martin P, Malirat N, Koval S, Raffel R, Sobhian B, Severac D, Rialle S et al. 2014. Integrator complex regulates NELF-mediated RNA polymerase II pause/release and processivity at coding genes. Nature communications 5: 5531.

Sulkowska JI, Morcos F, Weigt M, Hwa T, Onuchic JN. 2012. Genomics-aided structure prediction. Proceedings of the National Academy of Sciences of the United States of America 109: 10340–10345.

Sullivan KD, Steiniger M, Marzluff WF. 2009. A core complex of CPSF73, CPSF100, and Symplekin may form two different cleavage factors for processing of poly(A) and histone mRNAs. Molecular cell 34: 322–332.

Teppa E, Zea DJ, Marino-Buslje C. 2017. Protein-protein interactions leave evolutionary footprints: High molecular coevolution at the core of interfaces. Protein Sci 26: 2438–2444.

Tesileanu T, Colwell LJ, Leibler S. 2015. Protein Sectors: Statistical Coupling Analysis versus Conservation. Plos Computational Biology 11.

Thakur TS, Dubey R, Desiraju GR. 2015. Crystal structure and prediction. Annu Rev Phys Chem 66: 21–42.

Uguzzoni G, John Lovis S, Oteri F, Schug A, Szurmant H, Weigt M. 2017. Large-scale identification of coevolution signals across homo-oligomeric protein interfaces by direct coupling analysis. Proceedings of the National Academy of Sciences of the United States of America 114: E2662–E2671.

Wang J, Mao K, Zhao Y, Zeng C, Xiang J, Zhang Y, Xiao Y. 2017. Optimization of RNA 3D structure prediction using evolutionary restraints of nucleotide-nucleotide interactions from direct coupling analysis. Nucleic Acids Res 45: 6299–6309.

Wu Y, Albrecht TR, Baillat D, Wagner EJ, Tong L. 2017. Molecular basis for the interaction between Integrator subunits IntS9 and IntS11 and its functional importance. Proceedings of the National Academy of Sciences of the United States of America 114: 4394–4399.

Xue LC, Dobbs D, Bonvin AM, Honavar V. 2015. Computational prediction of protein interfaces: A review of data driven methods. FEBS Lett 589: 3516–3526.

Yu JC, Vavrusa M, Andreani J, Rey J, Tuffery P, Guerois R. 2016. InterEvDock: a docking server to predict the structure of protein-protein interactions using evolutionary information. Nucleic Acids Research 44: W542–W549.

